# BTEXgenie: A curated and user-friendly tool for profile HMM-based substrate-specific annotation of BTEX degradation genes

**DOI:** 10.64898/2026.05.12.724592

**Authors:** June Qu, Arkadiy I. Garber, Catherine R. Armbruster

## Abstract

**Background:** Benzene, toluene, ethylbenzene, and xylene (BTEX) are volatile aromatic hydrocarbons that are widespread environmental pollutants arising from petroleum processing, fuel combustion, and other industrial activities. Persistent BTEX contamination poses substantial risks to human health and ecosystems, underscoring the need for effective long term remediation strategies. Microbial bioremediation is a promising and sustainable approach for BTEX removal, but development of these approaches requires accurate detection of the genes and pathways responsible for substrate specific degradation. Although profile hidden Markov model (HMM) databases are widely used for functional annotation, existing annotation resources lack the substrate-specific resolution needed to distinguish between closely-related BTEX-degrading enzymes with different catalytic specificities.

**Results:** We developed BTEXgenie as a sensitive annotation tool that uses custom HMMs built from alignments of experimentally validated BTEX degradation proteins to identify genes involved in the initial steps of aerobic and anaerobic BTEX degradation. BTEXgenie improved detection of anaerobic BTEX degradation genes that were absent from KOfam annotations. In benchmarking against the KEGG KOfam HMM database, BTEXgenie achieved 17.73%higher overall sensitivity while maintaining comparable specificity at 97.02%across genes involved in BTEX degradation pathways.

When applied to environmental metagenomes, BTEXgenie recovered pathway patterns consistent with reported site characteristics and known degradation potential. In addition to gene annotation, BTEXgenie supports downstream interpretation through KEGG pathway-based visualization of detected functions and Circos-based visualization of genomic hit distributions.

**Conclusions:** BTEXgenie is a substrate-specific annotation tool built from custom HMMs for detecting genes involved in BTEX degradation. By integrating gene annotation with pathway and genome-level visualizations, BTEXgenie facilitates characterization of microbial BTEX degradation potential in environmental and comparative genomic studies.

## Background

Environmental contamination arising from steel production, petroleum refining, and other industrial activities is difficult to remediate and poses persistent risks to environmental and human health. Among these pollutants, benzene, toluene, ethylbenzene, and xylenes (BTEX) are volatile aromatic hydrocarbons that are widespread in contaminated soils, sediments, and groundwater [1]. Long term exposure to BTEX is associated with adverse health effects, including neurotoxicity, and BTEX compounds are included on the Agency for Toxic Substances and Disease Registry (ATSDR) Substance Priority List [2, 3]. Microbial bioremediation offers a sustainable and cost effective approach by using the natural metabolic activities of bacteria to transform BTEX compounds to non-carcinogenic byproducts [4, 5]. Thus, accurate detection of BTEX degrading genes is important for identifying candidate degraders, assessing the metabolic potential of environmental communities, and monitoring remediation progress.

A common approach for functional gene annotation relies on BLASTp to compare protein sequences against reference databases [6], but pairwise similarity often fails to distinguish closely related enzymes with different substrates or catalytic properties. In comparison, profile Hidden Markov Models (HMMs) improve sensitivity by capturing conserved patterns from multiple sequence alignments, and resources such as Pfam, KOfam, and eggNOG HMM databases are widely used for protein annotation [7–9]. Yet, because these databases are designed for broad functional coverage, they often lack the resolution needed to identify specialized BTEX-degrading enzymes. More targeted resources such as CANT-HYD, HADEG, AnHyDeg, and AromaDeg have helped address this gap through profile HMMs, curated reference sequences, or phylogenomic placement [10–13], but distinguishing enzymes that differ in substrate specificity or regioselectivity remains difficult. This is particularly important for BTEX degradation, where, for example, aerobic toluene degradation can proceed through either monooxygenase or dioxygenase pathways, and different toluene monooxygenases hydroxylate distinct positions on the ring, leading to variable downstream products and degradation efficiencies [14].

Here, we present BTEXgenie, a functional annotation tool comprising 30 curated HMMs derived from experimentally validated enzymes involved in BTEX degradation (Figure 1). To improve sensitivity while preserving functional specificity, BTEXgenie incorporates iterative homolog expansion using jackhmmer from HMMER v.3.4 to retrieve distantly related homologs from UniProt [15, 16]. BTEXgenie focuses on enzymes involved in the initial activation steps of BTEX degradation, because these enzymes are specific functional markers of an organism’s or community’s potential to degrade BTEX.

**Fig. 1:**
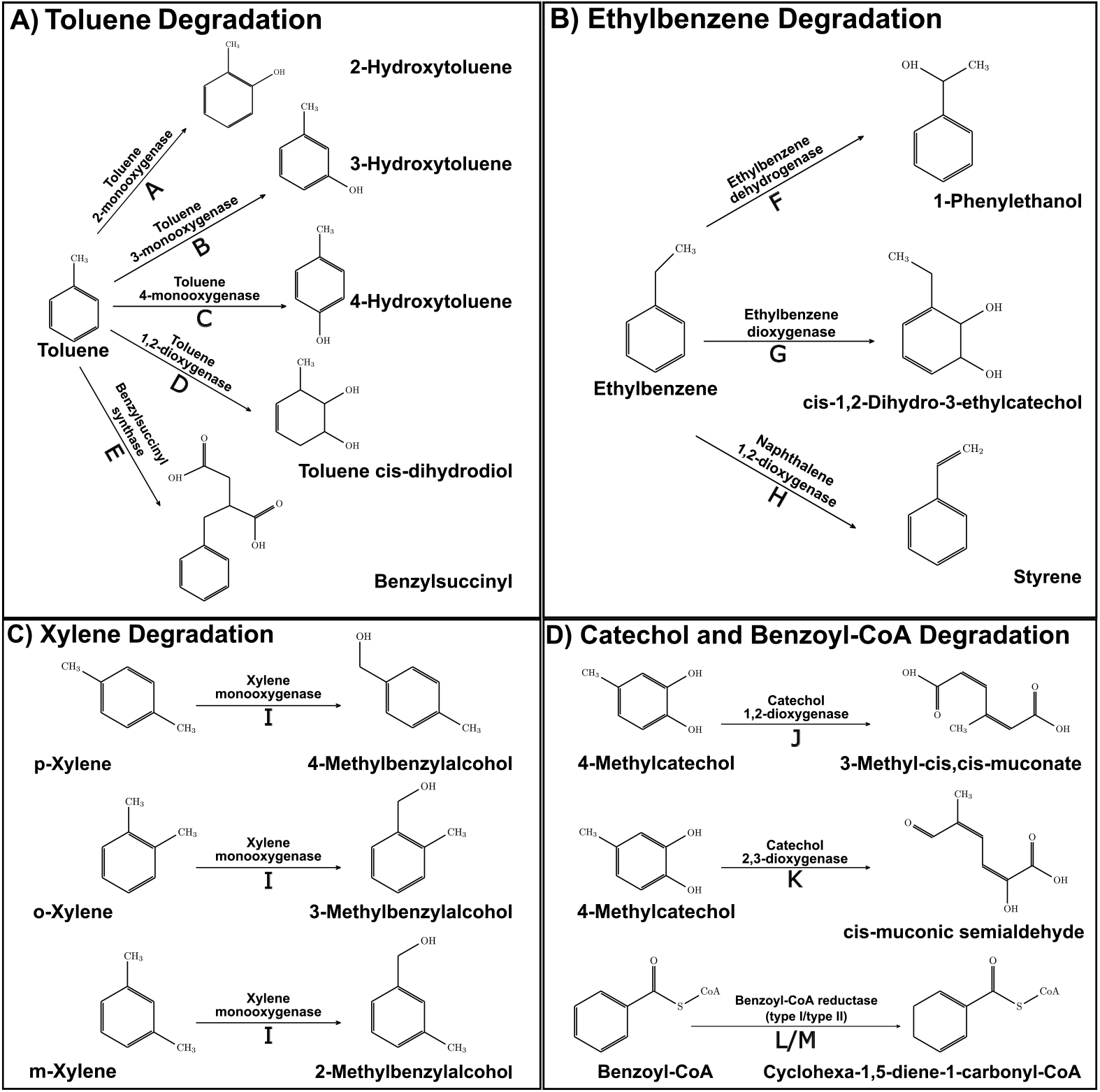
BTEX degradation reactions represented by BTEXgenie. Shown are the initial aerobic and anaerobic degradation steps for benzene and toluene (A), ethylbenzene (B), xylene (C), and reactions involved in processing central intermediates (D). Letters A to M denote enzymes represented by BTEXgenie HMMs.

## Implementation

### Pipeline Overview

- BTEXgenie scans input sequences against 30 curated profile HMMs representing enzymes that initiate BTEX degradation pathways.
- Users can provide either genome sequences, for which the tool performs gene predic-
- tion using Prodigal v2.6.3, or Prodigal predicted protein sequences as input;these are then screened against the BTEXgenie database and, optionally, the KOfam HMM database.
- Putative BTEX-degrading genes are filtered using model-specific thresholds and summarized in output CSV files.
- The pipeline generates two CSV files reporting detected BTEXgenie hits, a set of HTML files for pathway visualization, and a Circos plot for visualizing the distribution of hits on an isolate or metagenome-assembled genome (Table 1).

**Table 1:**
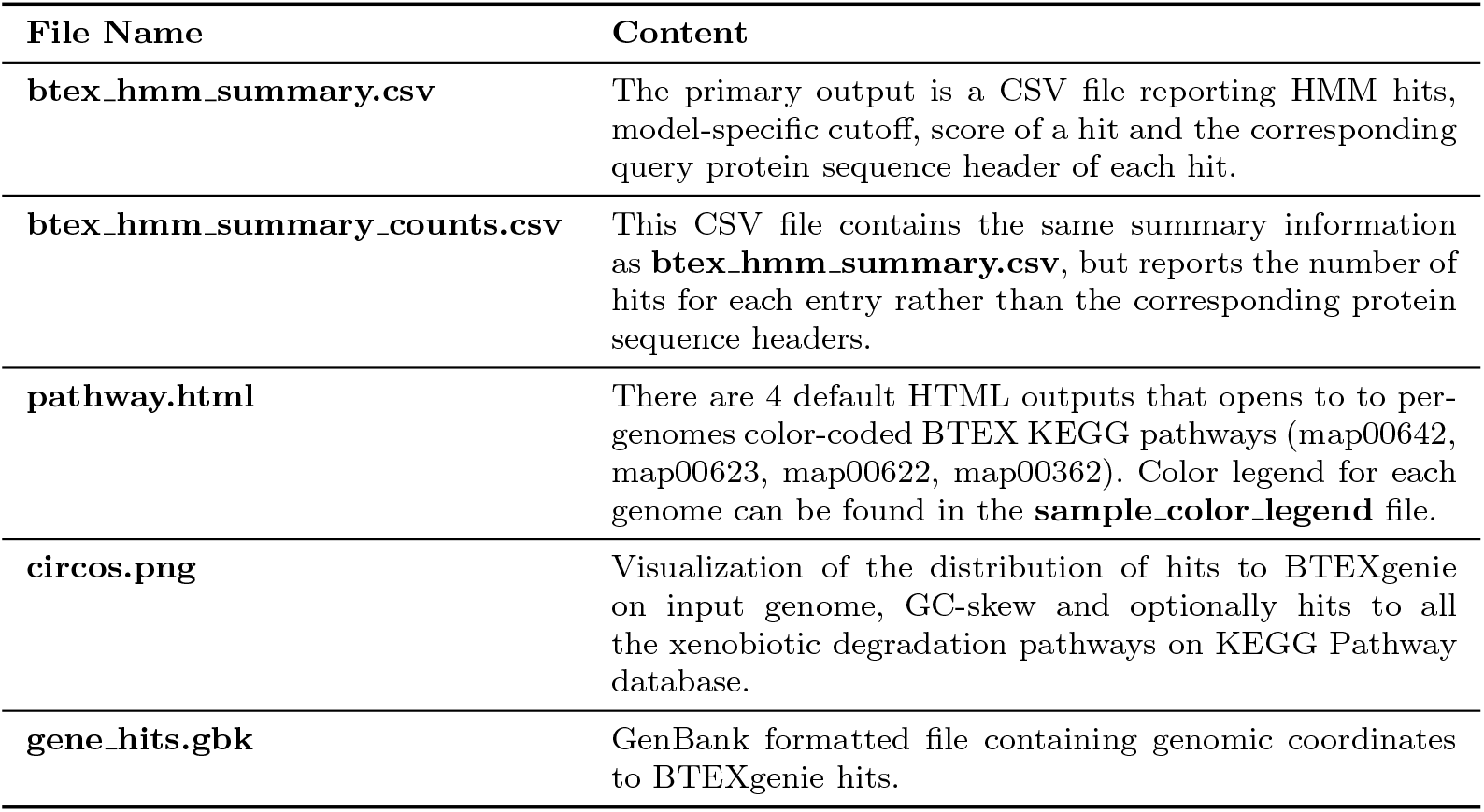
Main output files generated by BTEXgenie. Example visualization outputs (pathway.html and circos.png) are provided in Supplemental file 5.

### HMM Development

Protein sequences were collected separately for each target enzyme represented in Figure 1. For each enzyme, an initial high confidence positive set was manually curated from the literature and public databases and consisted of full length protein sequences supported by experimental evidence for the relevant biochemical activity and substrate specificity (Supplemental file 1). A corresponding negative set was assembled from homologous enzymes with related catalytic roles but different substrate specificities, representing likely sources of false positive matches.

### HMM threshold calibration

For each target enzyme, curated positive sequences were first deduplicated and clustered at 95%identity using USEARCH v11.0.667 i86linux32 [17]. Representative sequences were then aligned with MUSCLE v5.2, and alignment columns with greater than 50%gaps were trimmed using `esl-alimask` from HMMER v3.4 in majority mode [15, 18]. The trimmed alignment was used to build the initial profile hidden Markov model with `hmmbuild` from HMMER v3.4 [15]. This initial HMM was then evaluated by scanning both the curated positive and curated negative sequence sets to determine an initial detection threshold.

To improve the robustness of threshold selection, multiple candidate score thresholds were evaluated. For each candidate threshold, sequences scoring at or above the threshold were classified as positive, whereas those scoring below the threshold were classified as negative. This yielded counts of true positives, false positives, true negatives, and false negatives for each threshold. Threshold performance was then assessed using Matthews correlation coefficient, F1 score, and Youden’s J statistic:

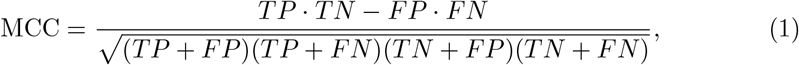

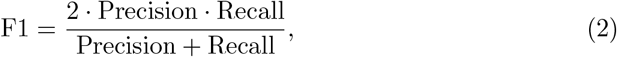

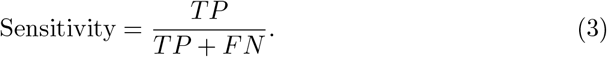

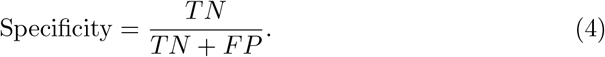

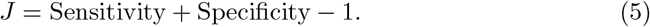

MCC was used as the primary performance metric because it incorporates all four outcome classes and remains informative under class imbalance [19]. F1 score was used to emphasize recovery of true positives while accounting for precision, and Youden’s J statistic was used to identify thresholds that balanced sensitivity and specificity [20, 21]. In addition to metric based threshold optimization, we also considered a gap criterion, in which the candidate threshold was placed at the midpoint of the largest score gap between adjacent sequences with different class labels [22]. This criterion favored cutoffs positioned between positive and negative score distributions rather than within dense regions of a single class.

Furthermore, to reduce overfitting during threshold selection, cross validation was performed on the curated positive and curated negative sets. Within each fold, candidate thresholds were proposed using only the training partition according to each thresholding policy. Each candidate threshold was then evaluated on the held out test partition using MCC. For each enzyme model, the final thresholding policy was selected as the one achieving the highest mean test MCC across folds, which will prioritize generalization performance over optimistic predictions on training data. For enzymes with insufficient numbers of experimentally validated sequences to support cross validation (Supplemental file 1), the threshold was selected directly from the candidate policies using the entire curated positive and negative sets to selected for a threshold that maximizes MCC. These models are therefore supported by limited calibration evidence and should be interpreted alongside complementary evidence, including co-occurring enzyme signals and taxonomic context.

### HMM Refinement

To better capture sequence diversity within each enzyme family, HMMs were refined through iterative homolog expansion using `jackhmmer` from HMMER v.3.4 [15]. For each target enzyme, the initial HMM and its corresponding initial threshold were used to search UniProt iteratively for additional homologous sequences. Searches were run until convergence, with a maximum of five iterations. Retrieved sequences scoring above the initial threshold were retained as putative homologs and incorporated into the model refinement process.

A final HMM was then constructed from the combined set of curated positive sequences and accepted putative homologs. The detection threshold for the final model was recalibrated by scanning the refined HMM against both the curated negative set and the expanded homolog set. The final cutoff was selected using the same thresholding framework described above.

In some enzyme families, experimentally validated proteins with closely related substrates produced overlapping score distributions that could not be reliably separated into substrate-specific models. In these cases, sequences were combined into broader functional models representing shared substrate classes. For example, toluene and benzene dioxygenases were merged into a single model representing the potential for toluene or benzene dioxygenase-mediated oxidation. This approach was applied only when an enzyme model retrieved experimentally supported homologs associated with closely related substrates at comparable scores, and when the proteins were biologically plausible members of an overlapping functional class.

### BTEXgenie Usage and Outputs

BTEXgenie take protein sequence files as input and scans them against 30 profile HMMs representing enzymes involved in the initiation of BTEX degradation. Briefly, the `btex-annotate` command annotates input genomic data using these profile HMMs, with an optional scan against the KOfam HMM database to assess degradation and metabolic potential across all KEGG pathways. The `btex-vis` command is useful for users interested in visualizing per sample contributions across an entire KEGG pathway. Finally, the `btex-run-circos` command generates a Circos plot showing the genomic distribution of all hits detected by BTEXgenie, with an option to also display hits assigned to xenobiotic degradation pathways in the KEGG Pathway database. A summary of BTEXgenie outputs is provided in Table 1, and detailed usage instructions with examples are available at https://github.com/armbrusterlab/BTEXgenie.

## Results and discussion

### Performance evaluation highlights the utility of BTEXgenie for detecting anaerobic degradation genes

Using 56 bacterial genomes with experimental evidence for BTEX degradation, we benchmarked BTEXgenie against KOfam, Pfam, and eggNOG to evaluate detection of BTEX degradation genes (Supplemental files 2 and 3). Because KOfam provides the closest annotation counterparts to BTEXgenie HMM targets, we directly compared the two tools using sensitivity and specificity calculated from reported gene presence and absence across 30 BTEX degradation genes (Figure 2).

**Fig. 2:**
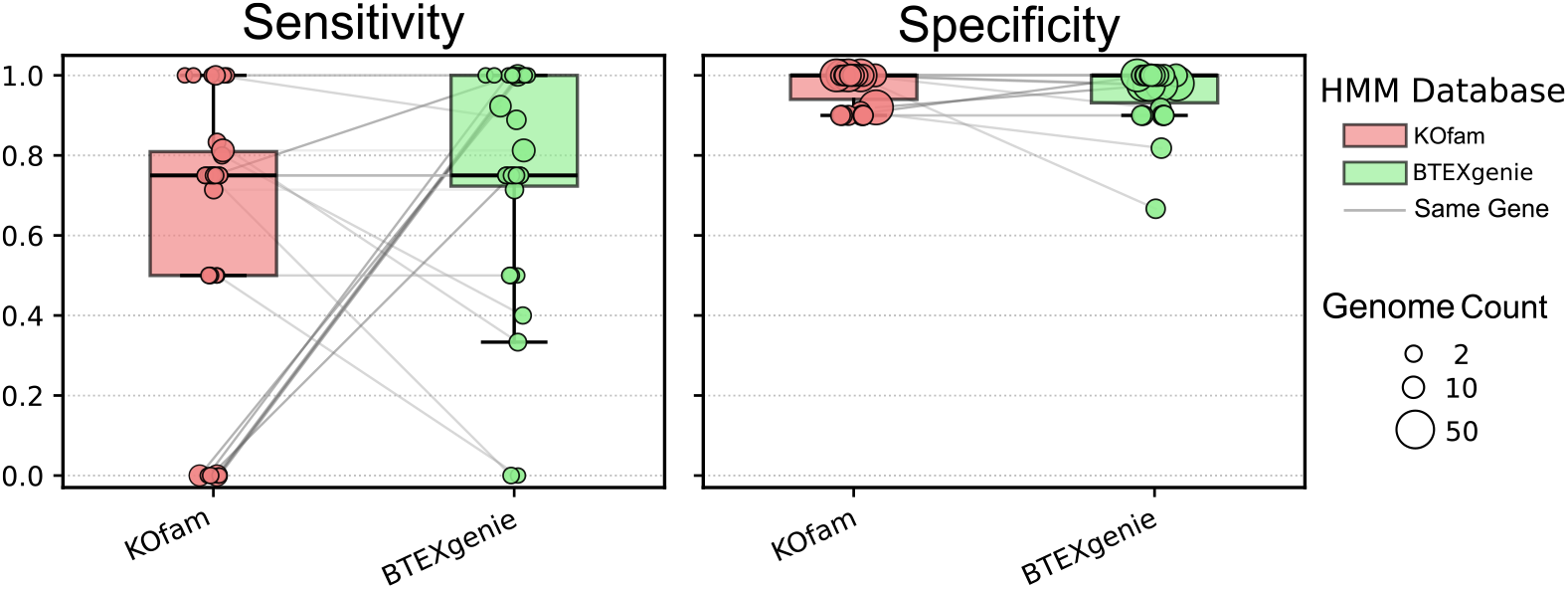
Comparison of KOfam and BTEXgenie HMM sensitivity and specificity for experimentally validated BTEX degradation genes. Each point represents the sensitivity or specificity calculated from a HMM from either KOfam (red) or BTEXgenie (green). Metrics are calculated using gene presence or absence for a set of experimentally validated genomes. Point size indicates the number of genomes contributing to each estimate, and gray lines connect the equivalent HMMs represented by KOfam and BTEXgenie.

Per-gene specificity was comparable between BTEXgenie and KOfam across all 30 genes, while sensitivity varied by target. Of the 30 genes, 17 showed equivalent sensitivity between tools, 5 showed lower sensitivity with BTEXgenie, and 8 showed higher sensitivity with BTEXgenie (Figure 2). Sensitivity gains were most pronounced for anaerobic BTEX degradation HMMs, particularly those targeting benzylsuccinate synthase (Bss) and ethylbenzene dehydrogenase (Ebd). Among 13 genomes with experimental evidence for anaerobic degradation via the Bss pathway, BTEXgenie produced a single false negative (*bssB* in *Desulfoscipio gibsoniae*), whereas KOfam failed to detect any Bss genes in these genomes (Figure 3). BTEXgenie also correctly identified Ebd in *Aromatoleum bremense* PbN1T, which KOfam did not detect [23].

**Fig. 3:**
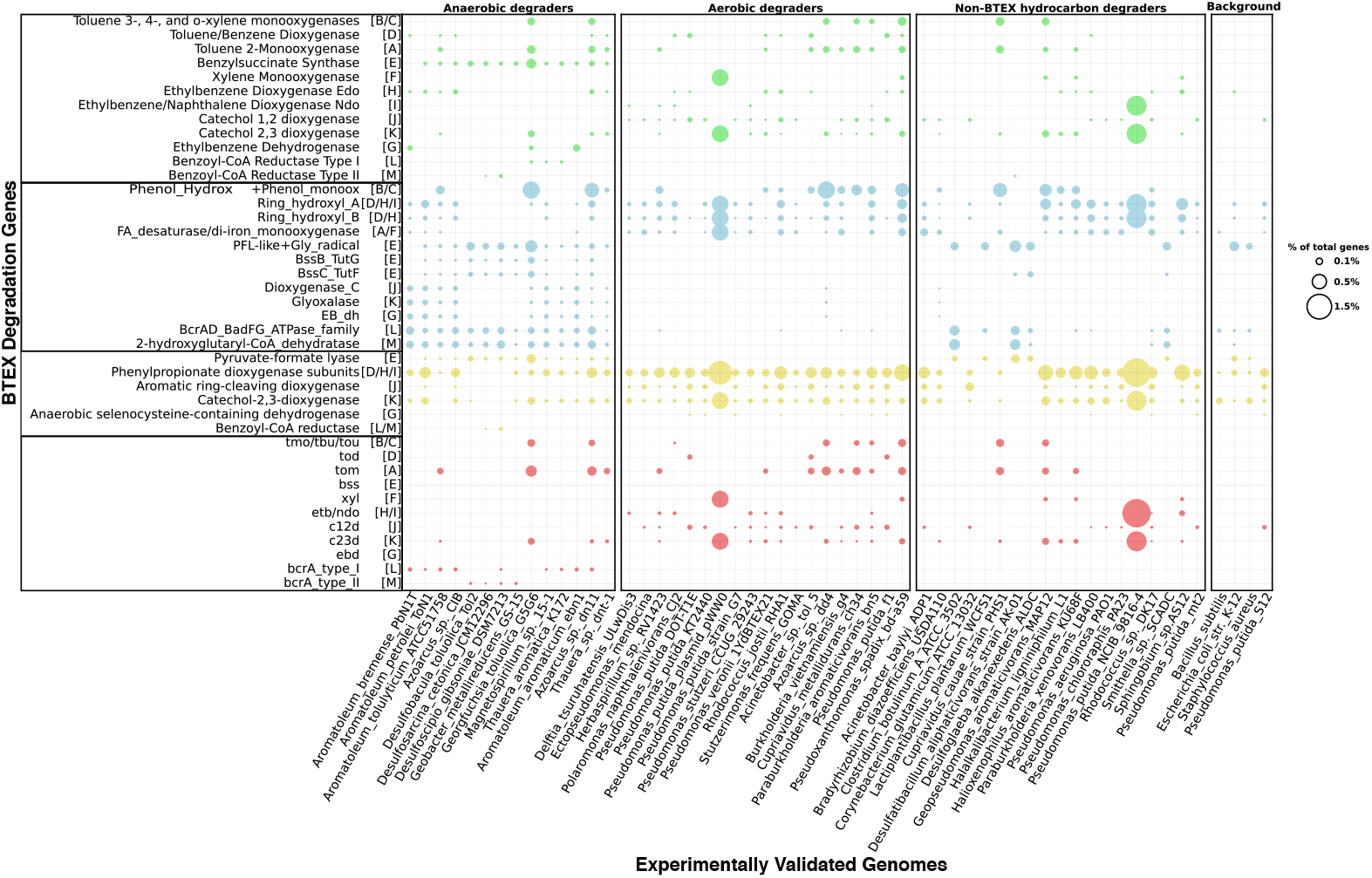
Validation of BTEXgenie against KOfam, Pfam, and eggNOG across 56 experimentally validated isolate genomes. Genomes are grouped into anaerobic BTEX degraders, aerobic BTEX degraders, non-BTEX hydrocarbon degraders and background genomes with no known BTEX degradation capability. Dot size represents the percentage of hits for each gene normalized to the total number of genes predicted in the genome. Dot color indicates the database used for annotation: BTEXgenie (green), Pfam (blue), eggNOG (yellow), and KOfam (red) HMM databases. Bracketed letters indicate the BTEX-associated enzymes that catalyze the same reaction and correspond to the enzyme labels shown in Figure 1.

KOfam showed higher sensitivity than BTEXgenie for benzoyl-CoA reductase types I and II. However, these comparisons were supported by smaller validation sets (6 and 5 genomes, respectively) relative to the 13 genomes used to evaluate Bss genes, and should be interpreted accordingly. Accounting for differences in validation set size, BTEXgenie achieved a pooled sensitivity of 75.89%and specificity of 97.02%, compared to 58.16%sensitivity and 97.82%specificity for KOfam, with the difference attributable primarily to improved detection of anaerobic degradation genes.

Pfam and eggNOG annotations broadly recapitulated the distinction between anaerobic and aerobic degraders but lacked the specificity of BTEXgenie. Anaerobic degraders were enriched for annotations corresponding to benzylsuccinate synthase-associated glycyl radical chemistry (enzyme E), while aerobic degraders were enriched for aromatic ring-hydroxylating dioxygenase activity (enzymes D/H/I), patterns consistent with the known degradation potential of the validated species. However, Pfam and eggNOG also produced false-positive signals in background genomes with no reported BTEX degradation capability, including hits to hydroxylase- and dioxygenase-associated domains and orthologous groups (Figure 3). Together, these results demonstrate the improved sensitivity and specificity of BTEXgenie for annotating BTEX degradation potential in experimentally validated isolate genomes.

### BTEXgenie improves substrate-specific detection of BTEX-degradation genes over Pfam and eggNOG

We evaluated the functional resolution provided by BTEXgenie relative to Pfam and eggNOG databases using five experimentally validated species (Table 2). While Pfam and eggNOG are broadly useful resources for protein family and orthologous group annotation, they are not designed for BTEX-specific catabolism and therefore often lack the substrate-level resolution needed to distinguish BTEX-degrading enzymes from functionally related proteins with no known role in BTEX catabolism [24].

**Table 2:**
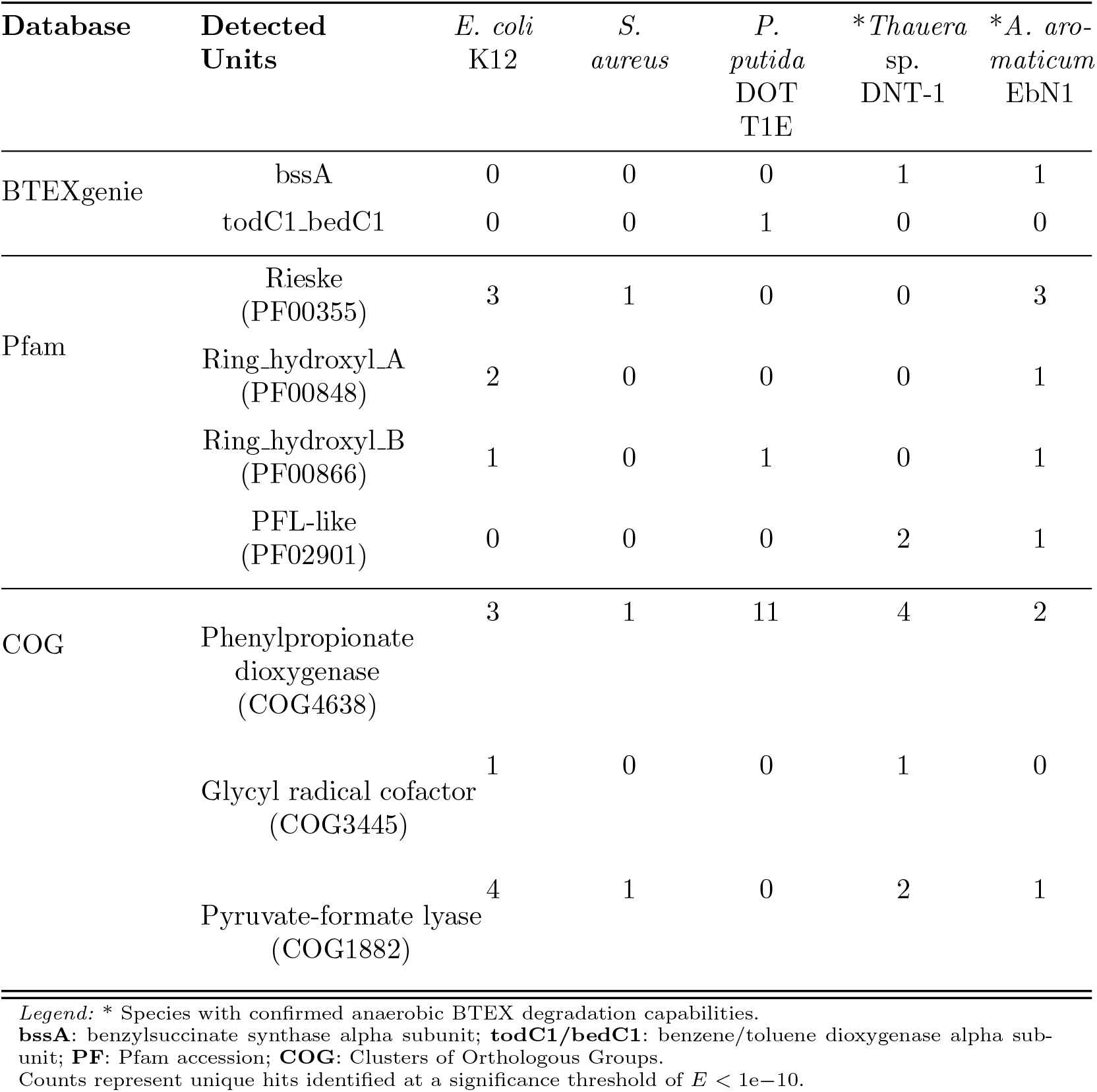
Gene hit counts across BTEXgenie, Pfam, and eggNOG databases.

For anaerobic BTEX degradation, Pfam detected pyruvate formate-lyase-like domain model (PF02901) in the confirmed anaerobic degraders *Thauera* sp. DNT-1 and *A. aromaticum* EbN1, and eggNOG returned broad orthologous group assignments including “glycyl radical cofactor”(COG3445) and “pyruvate formate-lyase”(COG1882) (Table 2). Although these annotations are consistent with the glycyl radical chemistry underlying benzylsuccinate synthase-mediated catalysis, they do not identify the BTEX-associated substrate or the specific metabolic reaction [25]. In contrast, BTEXgenie specifically detected the *bssA* subunit in both organisms using profile HMMs derived from experimentally validated protein subunits, providing direct evidence for the fumarate addition pathway responsible for alkylbenzene activation under anoxic conditions [26, 27].

Domain-level annotations can also produce misleading signals when used to infer BTEX-degradation ability. Toluene dioxygenase-related proteins belong to broad ring-hydroxylating dioxygenase families in Pfam that encompass enzymes involved in the metabolism of other hydrocarbons and structurally unrelated aromatic compounds. Consequently, Pfam can lead to detection of ring-hydroxylating dioxygenase domains in *E. coli* K12 and *S. aureus*, which has no reported BTEX-degrading ability [28]. Such false-positive signals may complicate assessment of BTEX-catabolism potential in genomic and metagenomic datasets.

For aerobic degradation, the closest Pfam and eggNOG counterparts to the BTEXgenie dioxygenase models are PF00866 and COG4638, both of which were detected in *P. putida* DOT-T1E. However, these annotations resolve only to the level of ring-hydroxylating dioxygenase activity and phenylpropionate-related reactions, and lack the substrate specificity required to confirm BTEX-associated catabolism. BTEXgenie addresses this limitation through HMMs such as *todC1 bedC1*, which directly annotates the aerobic toluene dioxygenase route and enables unambiguous classification of the initial oxidation step in toluene catabolism [14].

### Distinct BTEX degradation profile from BTEX-containing environments

We applied BTEXgenie to 16 environmental metagenomes from diverse settings, including petroleum contaminated landfills, soil, seawater, and cultures enriched with BTEX or other hydrocarbon compounds (Supplemental file 4). BTEXgenie distinguished BTEX-contaminated environments from non-contaminated environments based on the presence of genes involved in pathway initiation.

Across metagenomes from a contaminated site undergoing wastewater bioremediation [29], BTEXgenie recovered clear shifts in both the abundance and composition of BTEX-associated functions across remediation stages. In particular, BTEXgenie detected 18 proteins with BTEX-related functions in the metagenomic sample from site p4 in the Netherlands wastewater remediation system, where water was undergoing treatment, whereas no BTEX-related hits were detected at site p3, which had contaminated groundwater. Consistent with this pattern, Figure 4 shows that samples from sites p1 to p3 form a distinct cluster characterized by the absence of BTEX-related enzymes, while samples from sites p4 to p7 cluster separately and show stronger BTEX-related signals. This result agrees with the changes in contamination across stages of the bioremediation process as described in the original study [29]. In addition, the bioremediation strategy used an aerated bioreactor, indicating that aerobic degradation pathways should be the dominant mode of BTEX degradation in the treatment system. Consistent with this expectation, BTEXgenie detected only aerobic BTEX degradation pathways in samples from sites p4 to p7 and did not detect anaerobic pathways in these remediation-stage samples.

**Fig. 4:**
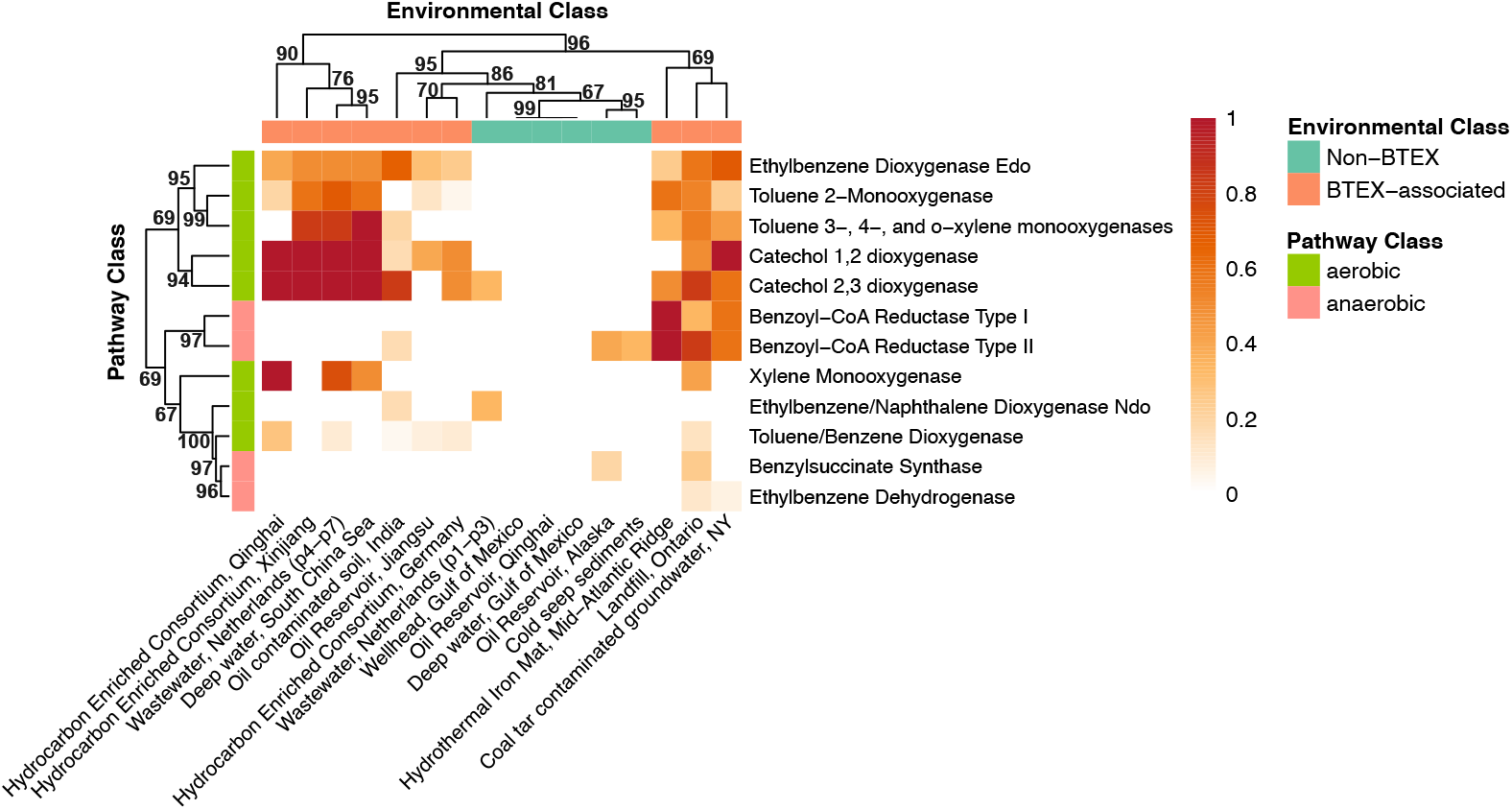
BTEXgenie hit profiles across 16 metagenomes from diverse environments and enrichment cultures. Color intensity reflects enzyme complex completeness, defined as the proportion of subunits detected within 15,000 bp of one another, with darker shading indicating more complete recovery of the corresponding pathway associated enzyme complex in a given sample. Metagenomes reported to contain BTEX compounds, or derived from BTEX or hydrocarbon enriched cultures, generally exhibited stronger and more complete pathway signals than metagenomes from environments not reported to be associated with BTEX contamination.

Furthermore, we found that metagenomes from non-BTEX associated environments generally lacked evidence of BTEX initiation pathways (Figure 4). For example, BTEXgenie detected benzoyl-CoA reductase in metagenomes from oil reservoir systems in the Alaskan North Slope and the Scotian Basin seabed. However, this enzyme is associated with downstream aromatic compound processing rather than the initial activation of BTEX compounds. In addition, the original studies of the Qinghai oil reservoir and Gulf of Mexico seawater samples did not report substantial BTEX contamination. These sites were instead described primarily as communities dominated by methanogenic and thermophilic microorganisms, rather than taxa typically associated with BTEX degradation [30, 31]. This interpretation is consistent with the lack of detected enzymes involved in the initial steps of BTEX degradation [32, 33]. In contrast, metagenomes from BTEX-associated environments, including the deep-water South China Sea (NCBI BioProject PRJNA1236222) and oil-contaminated soil from India [34], contained hits to enzymes representing both BTEX initiation steps and downstream intermediate-processing reactions . Across the 10 BTEX-associated metagenomes, we detected 673 hits assigned to BTEX initiation functions and 367 hits assigned to intermediate-processing functions. In comparison, among the 6 metagenomes from environments not reported to have substantial BTEX contamination, only 15 hits to BTEX initiation functions and 18 hits to intermediate-processing functions were detected.

Interestingly, hierarchical clustering based on the presence and absence of enzymes and completeness of enzyme unit separated the pathways into broad aerobic and anaerobic clusters. For example, benzylsuccinate synthase and ethylbenzene dehydrogenase, which are involved in anaerobic toluene and ethylbenzene degradation, clustered with benzoyl-CoA reductase rather than with most aerobic BTEX degradation enzymes. This pattern is biologically plausible because these enzymes participate together in the aromatic degradation processes [35]. While we did observe some aerobic functions associated with xylene and benzene degradation grouped within this cluster, the overall trend suggests co-occurrence of anaerobic degradation potential across these samples and is consistent with previous studies that use benzoyl-CoA reductase as a biomarker of anaerobic toluene degradation [36].

Together, these results indicate that BTEXgenie can accurately distinguish metagenomes enriched in genes for initial BTEX activation from metagenomes that contain only broader aromatic hydrocarbon processing capacity. BTEXgenie can also be applied to characterize anaerobic and aerobic BTEX degradation potentials in metagenomic datasets.

### Comparison of BTEXgenie to other hydrocarbon degradation enzyme databases

Existing hydrocarbon degradation databases, including HADEG and CANT-HYD, provide useful resources for annotating aerobic hydrocarbon degradation potential. However, in our analysis with BTEXgenie, these databases showed limited detection of anaerobic hydrocarbon degradation genes.

HADEG and CANT-HYD were developed for broad hydrocarbon degradation profiling, including alkanes and polyaromatic compounds, whereas BTEXgenie was developed specifically for BTEX degradation. To compare their applicability for BTEX focused analyses, we limited each database to genes associated with BTEX degradation. Only six genes were shared among all three databases, and all six were involved in aerobic toluene degradation. In contrast, nine genes were unique to BTEXgenie and were associated with anaerobic toluene and ethylbenzene degradation (Figure 5A). We further compared CANT-HYD and BTEXgenie using 10 experimentally validated genomes (Figure 5B). Both databases detected genes consistent with known aerobic and anaerobic BTEX degraders. However, CANT-HYD contained only eight aerobic degradation genes, while BTEXgenie contained 23, providing broader coverage of aerobic BTEX degradation potential. For anaerobic degradation, CANT-HYD included only BssA, whereas BTEXgenie covered seven genes, including subunits of benzylsuccinate synthase, ethylbenzene dehydrogenase, and benzoyl CoA reductase.

**Fig. 5:**
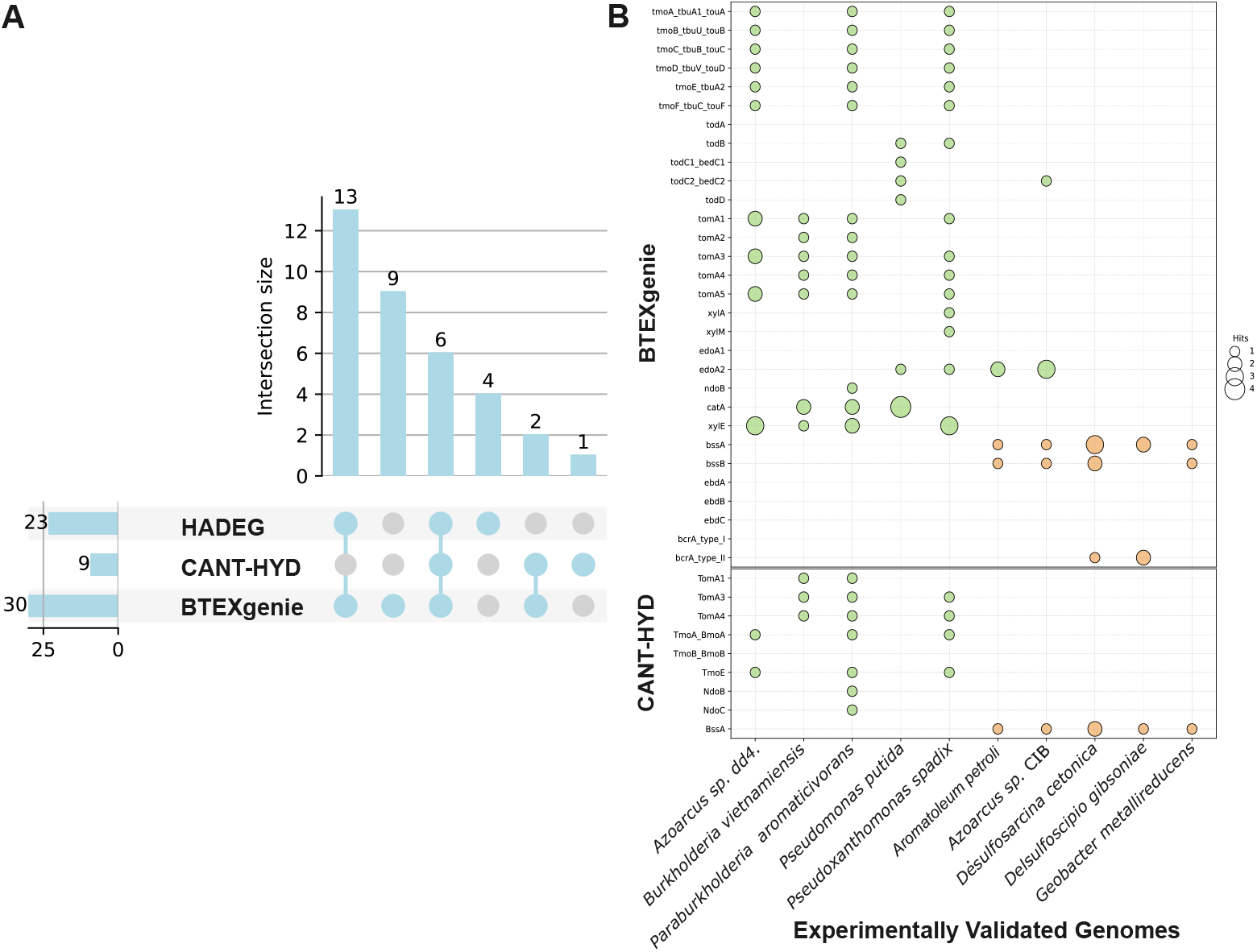
Comparison of BTEX degradation pathway annotations across 10 isolate genomes. These genomes were selected to highlight BTEX specific annotations provided by BTEXgenie relative to other hydrocarbon degradation databases. (A) UpSet plot showing pathway coverage shared among CANT-HYD, HADEG, and BTEXgenie, showing BTEXgenie has the most number of BTEX-specific annotations. (B) Comparison of gene hits identified by CANT-HYD and BTEXgenie. Dot size represents the number of hits for genes associated with aerobic pathways (green) and anaerobic pathways (orange).

Databases such as HADEG, AromaDeg, and HMDB support the prediction of genes involved in broad aerobic hydrocarbon degradation, but they do not capture genes associated with anaerobic degradation pathways. This limitation likely reflects the smaller number of experimentally characterized anaerobic degraders compared with aerobic systems. Therefore, the protein sequences used to build BTEXgenie represent the best currently available knowledge of anaerobic BTEX degradation and should be expanded as additional validated sequences are reported. Despite this limitation, our analysis shows that BTEXgenie is useful for providing an informative overview of BTEX degradation potential.

## Conclusions

We developed BTEXgenie as a user friendly tool with a curated profile HMM database designed specifically to detect and visualize enzymes that initiate aerobic and anaerobic BTEX degradation. The workflow is intended to be lightweight with an option to run additional HMM searches against the KOfam HMM database. It provides interpretable visualizations with KEGG pathways and Circos plots. In benchmarking against widely used annotation resources and analyses of environmental metagenomes, BTEXgenie recapitulated reported differences in degradation potential among environments while achieving improved sensitivity and specificity over KOfam annotations on experimentally validated genomes.

We note that for enzymes such as ethylbenzene dehydrogenase, only a small number of experimentally validated sequences are available. In these cases, the sequences used to build the corresponding HMMs overlapped with those that would otherwise serve as independent benchmarks, which can inflate performance. However, we retained these models to reflect the current state of knowledge and applied a conservative cutoff calibration using all available positives alongside curated negative and background sequence sets. As additional functional validations become available, these models and thresholds can be updated and strengthened.

## List of abbreviations

*BTEX:*: Benzene, toluene, ethylbenzene, and xylene
*COG:*: Clusters of Orthologous Groups
*HMM:*: Hidden Markov Model
*KO:*: KEGG Orthology
*Pfam:*: Protein families database

## Availability and requirements

- **Project name:** BTEXgenie: A curated and user-friendly tool for profile HMM-based substrate-specific annotation of BTEX degradation genes
- **Project home page:** https://github.com/armbrusterlab/BTEXgenie
- **Operating system(s):** Linux and macOS
- **Programming language:** Python, R
- **Other requirements:** Conda package manager
- **License:** AGPL-3.0 license
- **Any restrictions to use by non-academics:** None

## Declarations

### Ethics approval and consent to participate

Not applicable

### Consent for publication

Not applicable

### Availability of data and materials

The datasets generated and analyzed during the current study are available at https://github.com/armbrusterlab/BTEXgenie.

### Author Contributions

JQ carried out the sequence and literature searches, implemented BTEXgenie, performed all analyses, and drafted the manuscript. CA contributed to methodology development and supervised the work. AG contributed to software testing and refinement. All authors reviewed and approved the manuscript.

## Acknowledgements

We would like to thank Katherine Wang for testing the tool and providing valuable suggestions.

## Supplementary Information

**Supplemental file 1:** Table of accession IDs and references for all enzyme units used to build the initial HMMs.

**Supplemental file 2:** Mapping between enzyme name, gene name, Pfam ID, Pfam name, KEGG IDs, KEGG names and eggNOG categories.

**Supplemental file 3:** 56 Experimentally validated isolated genomes used to benchmark BTEXgenie, KOfam, Pfam and eggNOG HMM databases in Figure 3.

**Supplemental file 4:** 16 metagenomic studies with project ID and references used in Figure 4.

**Supplemental file 5:** Example outputs from btex-vis and btex-run-circos.

