## Supplemental file 5 for "BTEXgenie: A curated and user-friendly tool for profile HMM-based substrate-specific annotation of BTEX degradation genes"

### Supplemental 5

These are example visualization outputs from `btex-vis`, which maps detected hits onto BTEX-associated KEGG pathways, and from `btex-run-circos`, which visualizes KOfam and BTEXgenie hits in a Circos plot. Detailed usage and additional information are available at: `https://github.com/armbrusterlab/BTEXgenie`

Example `btex-vis` outputs (Benzoate, Toluene, Ethylbenzene and Xylene degradation pathways):

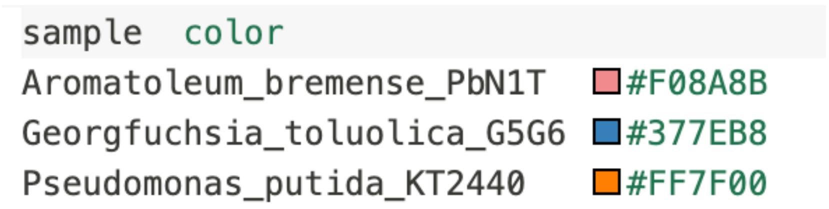

| sample | color |
| --- | --- |
| Aromatoleum_bremense_PbN1T | #F08A8B |
| Georgfuchsia_toluolica_G5G6 | #377EB8 |
| Pseudomonas_putida_KT2440 | #FF7F00 |

Figure 1: **Example `sample_color_legend` with per-sample colors used for annotating genes on KEGG pathways.**

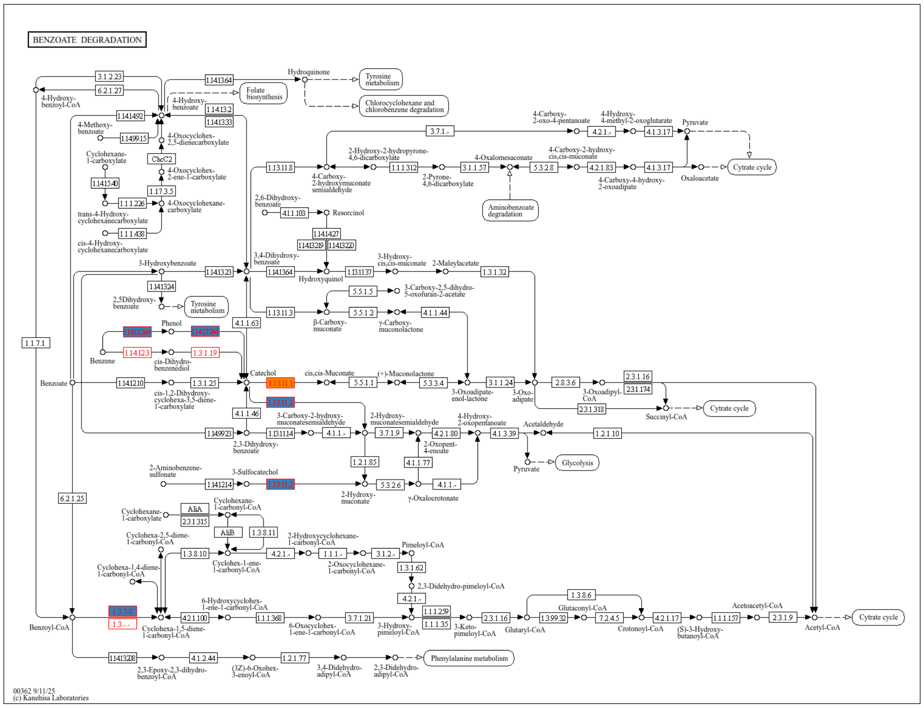


Figure 2: **Benzoate Degradation (map00362)**

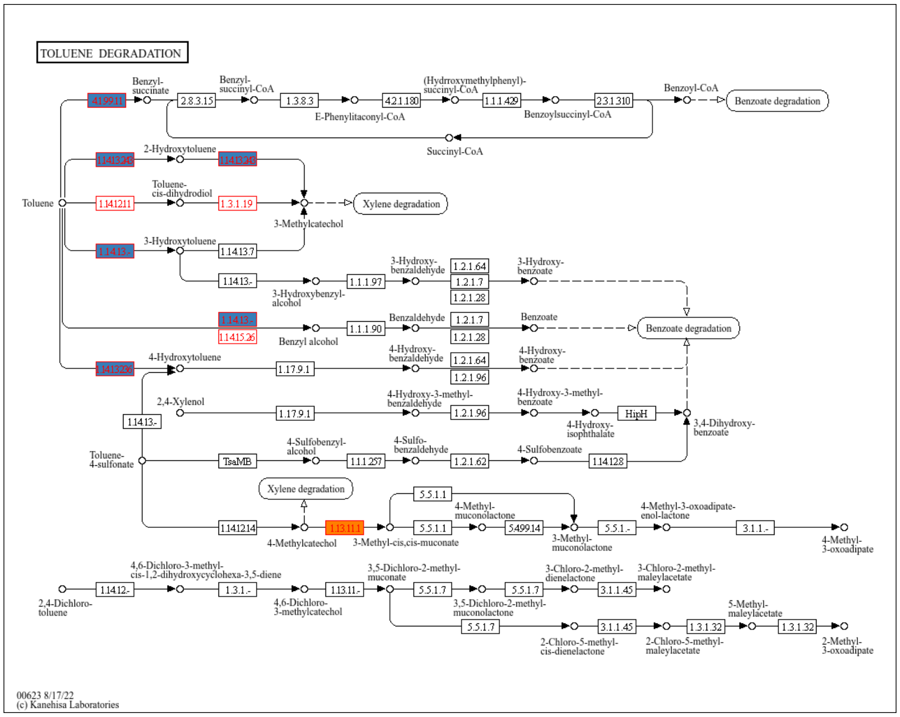


Figure 3: **Toluene Degradation (map00623)**

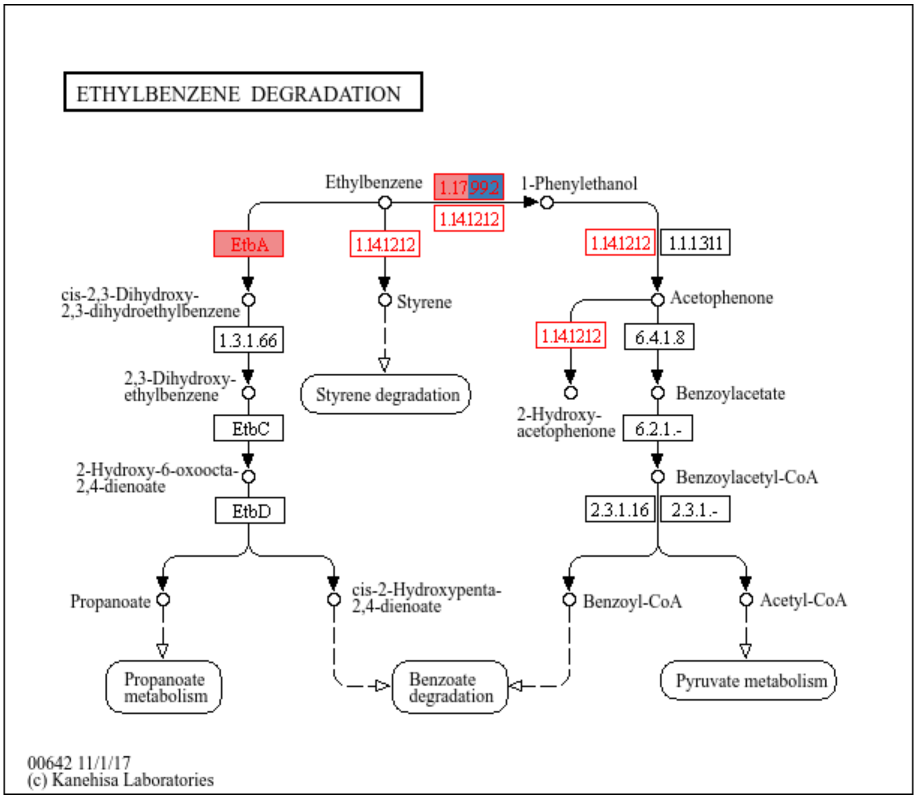


Figure 4: **Ethylbenzene Degradation (map00642)**

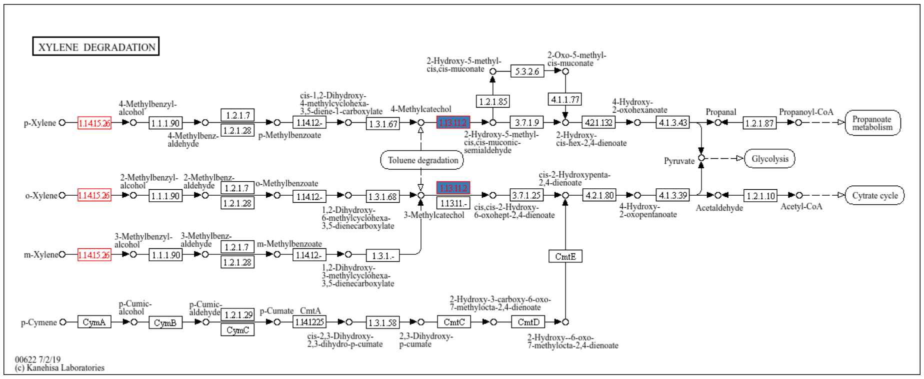


Figure 5: **Xylene Degradation (map00622)**

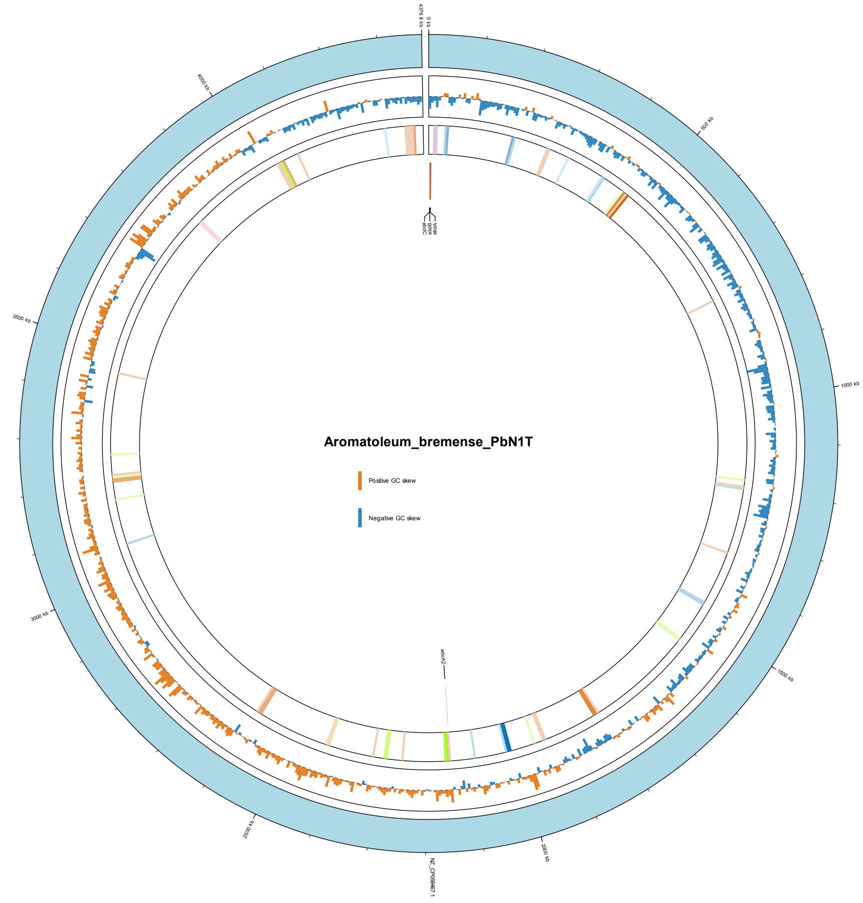


Figure 6: **Example output from `btex-run-circos` using the *Aromatoleum bremense* PbN1T genome.** From the outermost to innermost tracks, the plot shows the genome track, GC-skew track, hits to xenobiotic degradation pathways in KEGG, and BTEXgenie hits.
